# Proximity labelling allows to study novel factors in chloroplast development^a^

**DOI:** 10.1101/2022.12.08.519630

**Authors:** Bernhard Wurzinger, Simon Stael, Manuela Leonardelli, Carlo Perolo, Michael Melzer, Palak Chaturvedi, Leila Afjehi-Sadat, Wolfram Weckwerth, Markus Teige

## Abstract

Chloroplast development is initiated by light-signals triggering the expression of nuclear encoded chloroplast genes in a first phase, followed by massive structural changes in the transition from proplastids to mature chloroplasts in the second phase. While the molecular players involved in the first phase are currently emerging, regulatory components of the second phase, demanding high plastid translational capacity and RNA processing, are still enigmatic. This is mostly due to the very limited amount of plant material at the early phases of development that makes biochemical studies such as identifying protein interaction networks very difficult. To overcome this problem, we developed a TurboID-based proximity labelling workflow that requires only very limited sample amounts to obtain mechanistic insights into protein interaction networks present in the early stages of plastid development. We used the CGL20a protein, a novel factor involved in chloroplast development, as bait for *in vivo* proximity labelling in developing seedlings 7 days after germination. We found that CGL20a resides in a nexus of RNA binding proteins mainly associated to ribosomal RNA (rRNA) including different ribosome-associated proteins.

**One-sentence summary:** The use of plastid-specific in vivo proximity labelling in Arabidopsis seedlings allows to identify novel components in chloroplast development in higher plants.

## Introduction

Chloroplast development is the key to autotrophic life of all plants. As true organelles of endosymbiotic origin, chloroplasts develop from proplastids in response to light signals. This process involves two phases, a first phase where red-light signals trigger nuclear transcriptional responses (Waters and Langdale, 2009; Cackett et al., 2022), followed by internal retrograde chloroplast-to-nucleus signals for coordination of gene expression between nucleus- and plastid-encoded proteins (Dubreuil et al., 2018). The conversion of proplastids into chloroplasts is accompanied by high transcription levels of plastid- and nuclear-encoded genes involved in the transcription/translation apparatus, while genes required for photosynthesis are highly expressed only later in development. This leads then to massive structural changes in the chloroplast (Pipitone et al., 2021). These structural changes in the second phase require plastid transcription and translation and coordination with the nuclear-encoded components of the photosynthetic machinery after import into the plastids (Zoschke et al., 2018; Tadini et al., 2020). A recent imaging and biochemical analysis of 3-day-old Arabidopsis seedlings during de-etiolation revealed that within only 12 h of light, and continuing for about 4 days, a massive accumulation of proteins of mature photosynthetic complexes and expansion of the surface area of thylakoids can be observed (Pipitone et al., 2021). However, the regulation of this second phase is mechanistically not understood. During de-etiolation, massive protein synthesis is required for assembly of the highly abundant photosynthetic complexes embedded in thylakoids. Therefore, we selected the Arabidopsis CGL20 a/b proteins, which were recently described as factors affecting chloroplast development, particularly under cold conditions, by exerting influence on plastid ribosome processing (Reiter et al., 2020) to get further insights into processes involved in this second phase. The functionally redundant CGL20 a/b proteins are conserved in all *viridiplantae* and provide an interesting starting point for future studies in chloroplast development. In fractionation experiments from chloroplast stroma, CGL20a-GFP was found in high molecular weight fractions containing ribosomes, but in an RNA-independent manner (Reiter et al., 2020). This finding prompted us to test TurboID (TID)-based proximity labelling in plastids to identify interacting proteins.

So far, our mechanistic understanding of CGL20 functions has been derived from 6 – 12-week-old plants that developmentally resembled a green wild type phenotype. Here, we decided to focus on molecular interactors of CGL20a at an early stage of development, 7 days after germination, where we observed the biggest phenotypic differences between wild type and the *cgl20a/b* double knockout plants. Classical unbiased analysis techniques for plastid protein interactions in higher plants rely on plastid isolation and purification, rendering them suitable only for instances where plant material is not limiting. Additionally, robust methods to identify stable protein-protein interactions, such as affinity purification mass spectrometry (APMS), are limited in their ability to resolve transient interaction networks due to the low affinity between interaction partners and the high dilution occurring during protein purification. Therefore, we chose an *in vivo* proximity labelling approach to overcome these bottlenecks. TurboID based protein proximity labelling was recently successfully introduced into plant science (Mair et al., 2019; Arber et al., 2020). This system is based on a promiscuous biotin-ligase which essentially catalyses the formation of biotinoyl-5’-AMP which then is thought to be released and randomly attaching to primary amines within a radius 10-30 nM inside cellular compartments such as the cytosol (Branon et al., 2018; Roux et al., 2012; Choi-Rhee et al., 2004). However, there was concern if it would work in plastids of higher plants as no free biotin was detected in isolated chloroplasts, whereas ~11 μM of free biotin were reported for the cytosol (Badelt et al., 1993). Still, biotin-protein ligase activity was described in plastids (Tissot et al., 1998) as well as endogenous biotin-protein conjugates in plastids (Alban, 2011). We applied TID-based proximity labelling after complementing *cgl20a/b* null mutants with genomic fragments of CGL20a fused to TID under the control of a *CGL20a* 2kb promoter fragment. Suitable control lines enabled us to identify CGL20a spatio-temporally neighbouring proteins.

## Results

### Selection of suitable CGL20a-TID expressing *Arabidopsis* plant lines

Arabidopsis *cgl20 a/b* null mutants exhibit a strong virescent phenotype during the early stage of plant development (Figure 1A) but become phenotypically similar to Col0 wild type plants after ~ 8 weeks growth under low light conditions at temperatures between 18 and 26 °C. Still, also in 4-week-old leaves, the impaired plastid development is clearly visible in ultrastructural analysis (Figure 1B). EM pictures show a strongly reduced thylakoid stacking and swollen thylakoid membranes. *cgl20a/b* double knock-out plants were complemented with the genomic locus of *CGL20a* including a 2kb upstream fragment serving as promoter. A YFP-TID cassette replaced the *CGL20* STOP codon. A construct consisting of the same promoter/UTR and the CGL20a plastid-targeting peptide in exon1 fused to (YFP)-TID was used to create a control of free plastid stroma targeted TID in an *Arabidopsis Col0* background (Figure 1C). Segregation analysis from obtained transformant seeds was done to obtain single insert homozygous plants (Figure S1). It was previously shown that differential analysis of TID-based proximity labelling data is best interpretable if the bait-TID and control-TID constructs are similarly active (Mair et al., 2019). Therefore, in a first step we tested expression levels of the selected complemented lines by western blotting. Probing with a GFP antibody against the YFP tag in the respective TID constructs (Figure 1C) revealed that despite pheno-copying the Col0 phenotype in all lines, the expression levels of the respective TID constructs differed considerably between the lines (Figure S2A and B). In the control lines we observed one major band corresponding to the mature, ~65 kDa plastid-TID protein (Figure S2A). Interestingly, we always observed a double band at ~75 kDa and ~69 kDa for the fusion protein in all CGL20-TID lines, irrespective of its overall expression level (Figure S2B). This difference would roughly correspond to the size of the chloroplast targeting peptide (cTP) and may indicate that plastid import of CGL20a or cleavage of its transit peptide might be slow compared to the control despite using the identical cTP. Taking advantage of the YFP tag, we looked at the subcellular localisation of the different TID constructs by fluorescence microscopy and found that the CGL20a cTP does efficiently target the control TID into the plastid (Figure 1A). Also, for the CGL20a-TID construct we observed the YFP signal exclusively in plastids, confirming plastid localization of the mature CGL20a-TID protein (Figure 1A). Importantly, both lines resemble the wild type phenotype (Figure 1B). From this we concluded that the control construct has no obviously harmful effects on plant growth even after being propagated over 4 generations and if so, only minimally interferes with the endogenous biotinylation system (Tissot et al 1998). The phenotypic complementation of a *cgl20a/b* double knock out plant by the CGL20a-TID construct suggests that CGL20a is functional in its usual subcellular environment despite carrying the TID-YFP tag which is ~5 times bigger than the mature CGL20a protein. Finally, we selected two lines, designated as “cgl20a-TID L1” and “plastid-TID L1”, for our study based on the comparable levels of full length, mature TID constructs (Figure S2A and B).

**Figure 1.**
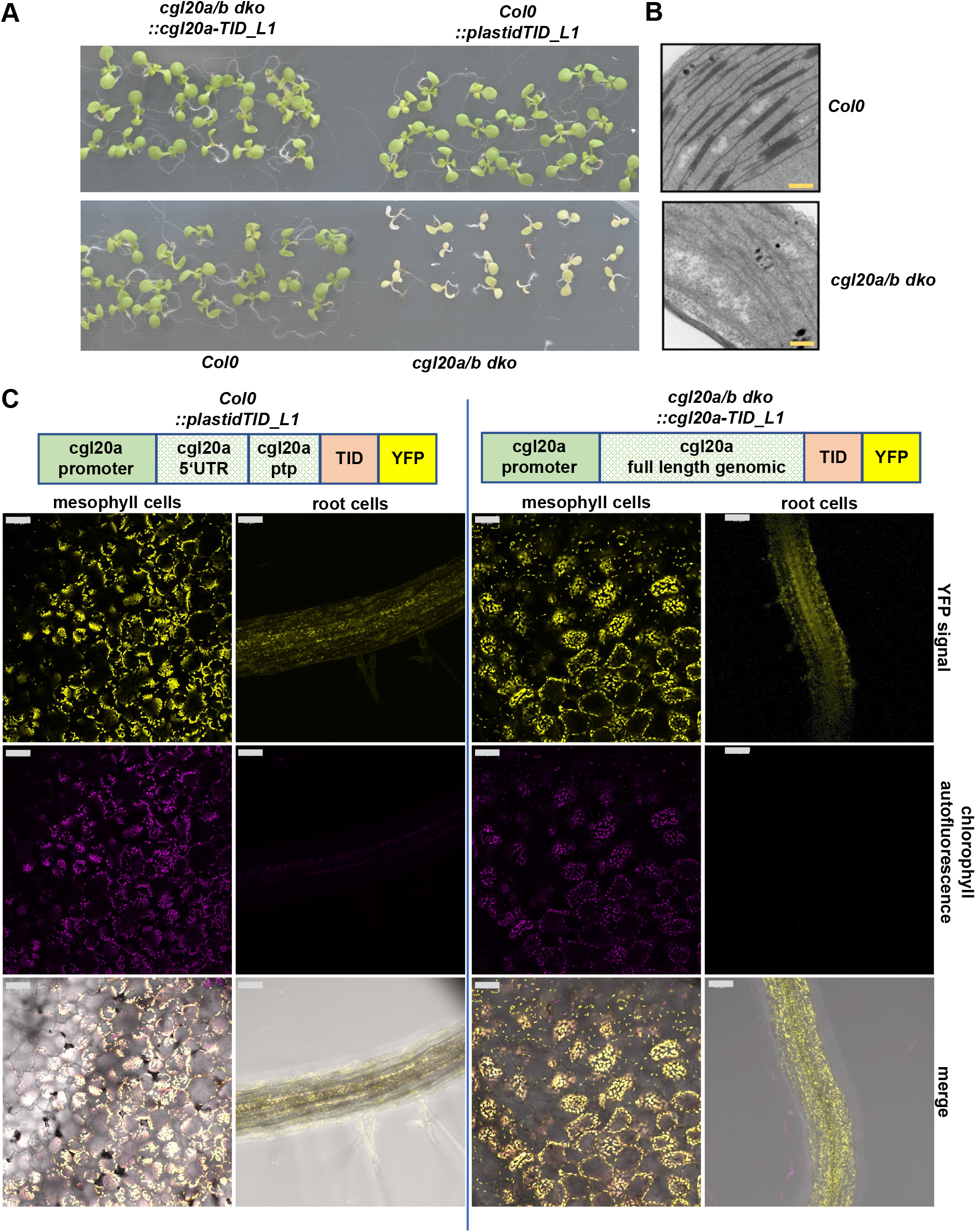
Constructs and lines used in this study. **A** shows phenotypes of *Col0* (lower left), *Col0::plastid TID* (upper left), *cgl20a/b dko* (lower right) and *cgl20a/b::cgl20a-TID* (upper right) seven day old seedlings. **B**, Transmission electron micrographs of representative *wild type Arabidopsis* chloroplasts (upper image) and *cgl20a/b dko* chloroplasts. Orange scale bars indicate 500 nm. **C**, schematic representation of constructs used in this study and sub cellular localisation of the plastid targeted TID-YFP and CGL20a-TID-YFP fusion proteins. The left panels show the "control” construct and its presence in leaf and root plastids, the right panels show the *cgl20a-TID-YFP* construct used for complementation in the *cgl20a/b dko* background. Grey scale bars represent 50 μm

### CGL20a-TID and plastid-TID show biotin ligase activity in seedlings

To our knowledge this is the first time that biotin ligase-based proximity labelling is applied inside plastids of higher plants. In plants, the main reactions for biotin synthesis occur in mitochondria. From there biotin finally needs to be translocated to the other cellular compartments such as the plastids where endogenous biotin ligase activity has been described (Alban et al., 2011). Surprisingly, it was reported that in plastids only protein bound biotin was detected but no free biotin, while concentrations of ~ 11 μM of free biotin were detected in the cytosol (Badelt et al., 1993). This gives rise to the question under which condition TID-based proximity labelling would take place in plastids? Therefore, we set up a test series treating plants with 50 μM exogenously applied biotin for different times (Figure 2) and subjected total plant protein extracts to western blot analysis with a biotin specific antibody. The plastid-TID L1 samples resembled almost the Col0 wild type biotinylation patterns without addition of extra biotin. Already after 30 minutes we observed strong auto-biotinylation of the plastid-TID control, which did further increase until 180 minutes whereas the biotinylation pattern of Col0 was similar to the control without biotin after 180 min of biotin treatment. (Figure 3). These results showed that exogenously supplied biotin is able to enter the plastid and leads to increased biotinylation activity of TID inside the plastid. Interestingly, for CGL20a-TID we observed an auto-biotinylation band without the addition of exogenous biotin. Still, a similar increase in autobiotinylation was visible for CGL20a-TID after 30 and 180 minutes of biotin pulse chase as seen for the plastid-TID control (Figure 3). Together, these experiments demonstrated that the activity of both plastid_TID and CGL20a-TID fusion proteins can be triggered by addition of exogenous biotin allowing for pulse chase experiments despite the fact that both constructs are stably present *in planta*.

**Figure 2.**
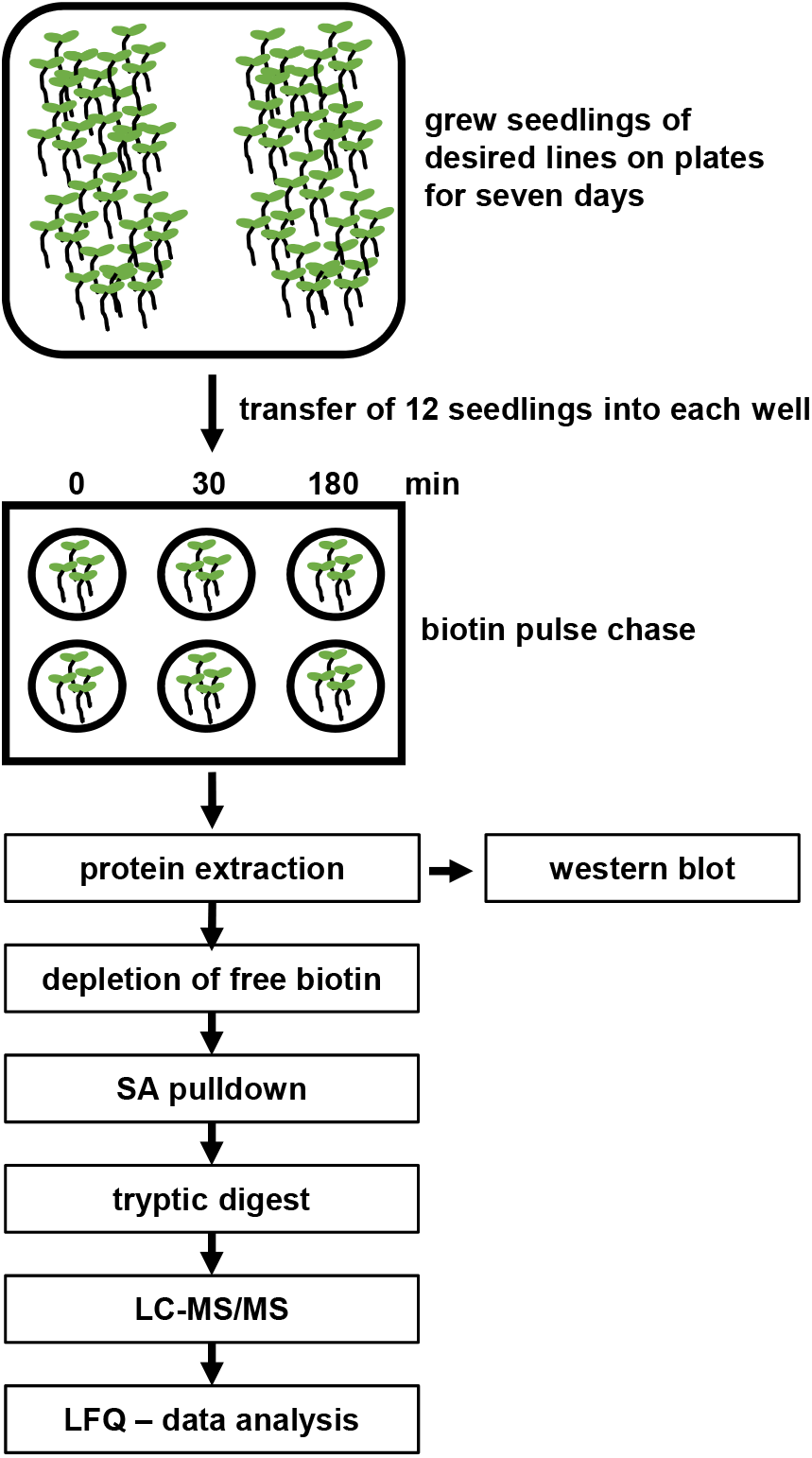
Scheme of work-flow. **A**, representative depiction of the experimental setup used for the biotin pulse chase experiment described in the text.

**Figure 3.**
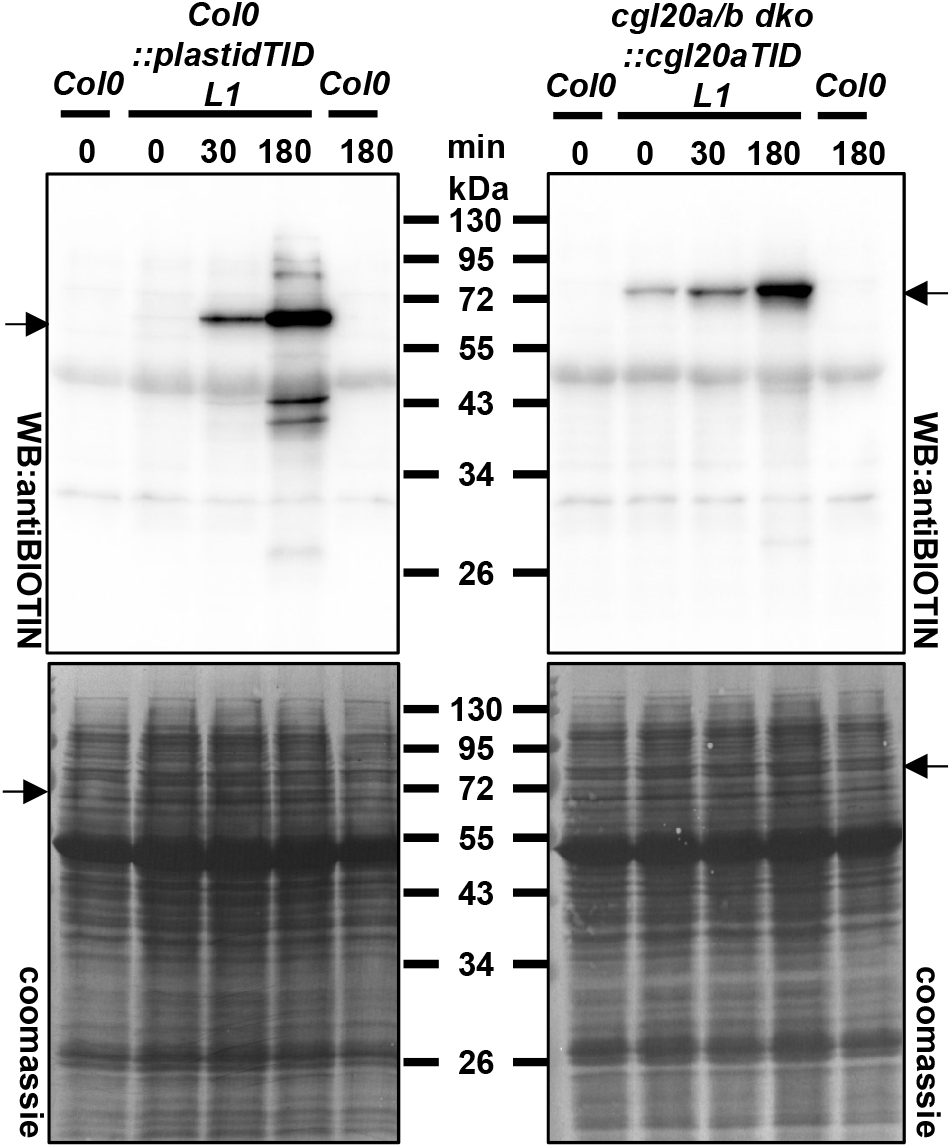
Biotin pulse chase western blot. The left panel shows the global biotinylome of a total protein extract obtained from *Col0::plastidTID L1* samples. The right panel shows the global biotinylome of a total protein extract obtained from *cgl20a/b dko::cgl20aTID L1* seedlings. The outermost left and right samples on each panel are extracts obtained from Col0 wild type seedlings serving as a null control for background biotinylation. The position of the respective full-length TID fusion proteins is indicated by arrows. In both cases protein extracts were separated on an SDS-PAGE gel and subseqeuntly analysed by western blotting with an antibody raised against biotin. The bottom images are the coomassie stained membranes after the respective western blots, serving as loading control.

### CGL20a-TID and plastid-TID predominantly biotinylate plastid proteins

Encouraged by the results described above we asked if the respective TID constructs are able to biotinylate plastid proteins. For this purpose, we chose to analyse the protein extracts of 12 seven-day old seedlings, per replicate, subjected to a 30 minutes biotin pulse chase, and compared labelling patterns to time point 0 (Figure 2). The 180 minutes timepoint was omitted from analysis since data presented by Mair and co-workers (Mair et al., 2019) indicated that after 3 hours of labelling saturation effects would take place, leading to an increased overlap of the control with the bait samples. Streptavidin (SA) beads were used to enrich biotinylated proteins from total protein extracts which were analysed by MS/MS after tryptic digest. We started our analysis from 1849 *Arabidopsis* protein groups identified throughout all samples (Table S1). From these we identified 708 protein groups enriched in the plastid-TID samples (Figure 4A) and 567 in the CGL20a-TID samples (Figure 4B) after the biotin pulse chase. Comparing these numbers to those of proteins enriched after 30 min biotin pulse chase and SA pulldown in Col0, 21 protein groups in total (Figure S3), we conclude that TID is responsible for more than 95 % of biotinylation events happening during the biotin pulse chase. Testing for the subcellular localisation of the enriched proteins by using SUBA5 (Hooper et al., 2017) we observe that 64 % of the 708 protein groups identified as enriched in the plastid-TID samples are reported to be plastid localised followed by 18 % in the cytoplasm and 6 % in mitochondria. In case of CGL20a-TID the numbers are similar, 62 % plastid localised, 21 % in the cytosol and 5 % in mitochondria. From these data we can conclude that indeed the majority of TID activity resides inside the plastid which is in good agreement with the YFP localisation data showing signal from plastids for the respective constructs (Figure 1A). Nevertheless, ~ 100 cytosolically annotated protein groups were found to be enriched in each of the TID lines which is approximately 5 times more than in the Col0 control which might be an indication that TID is partly active already as preprotein before being imported to the plastid.

**Figure 4.**
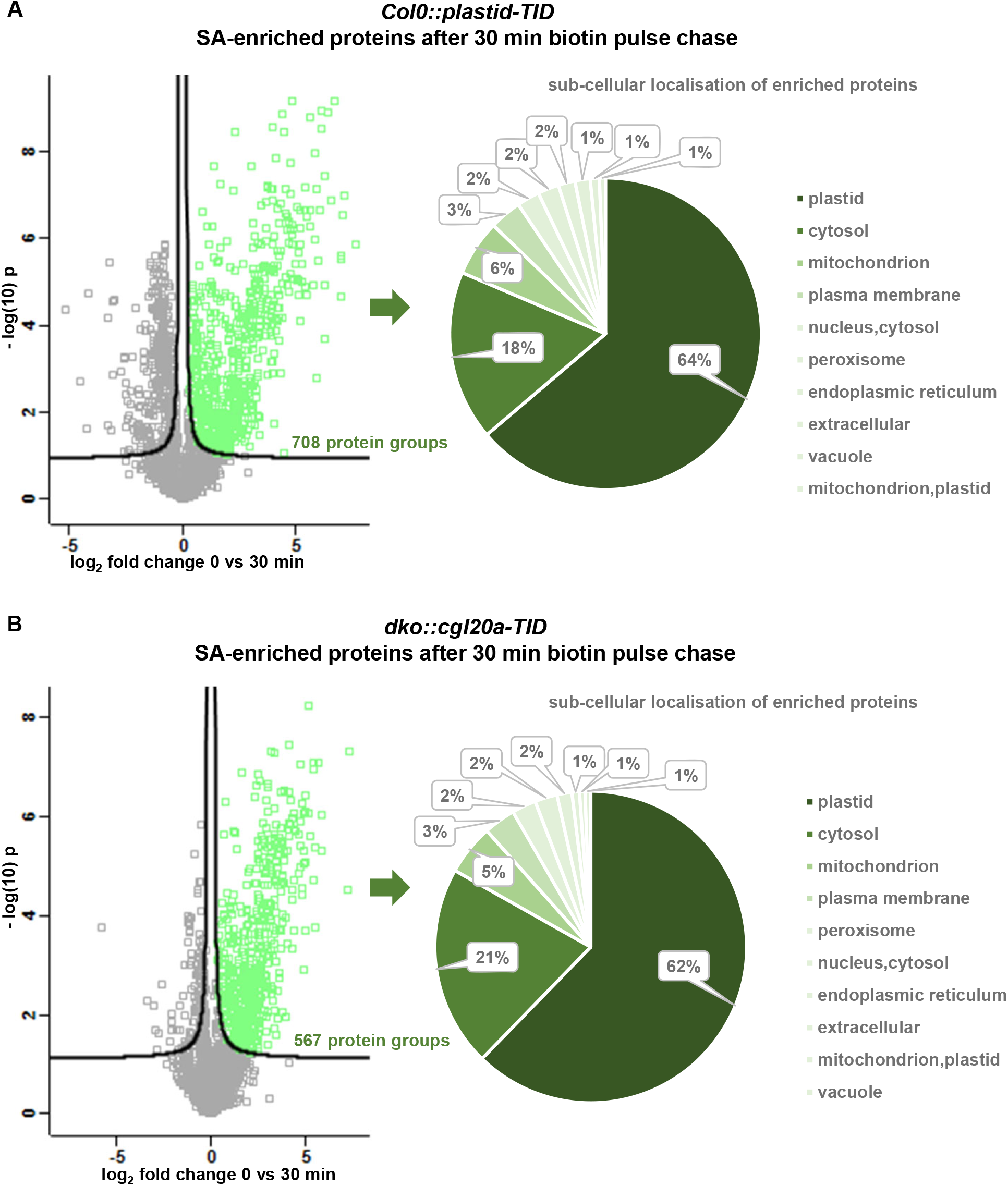
Biotin pulse chase enrichment analysis. **A**, left panel, volcano plot of SA enriched protein groups from *Col0::plastidTID* protein extracts. Green squares indicate significantly SA-enriched protein groups after 30 minutes of biotin pulse chase. The left panel indicates the sub-cellular distribution of all SA-enriched protein groups after 30 minutes of biotin pulse chase. **B**, similar to **A** for SA enriched proteins of *cgl20a/b dko::cgl20aTID* complementation line protein extracts. Sub-cellular localisation classification was done with SUBA5. Solid black lines indicate significance thresholds for an FDR of 0.05. 5 biological replicates were analysed for each genotype, condition and timepoint

### Proteins of the plastid import machinery are enriched in CGL20a-TID samples

Since both, CGL20a-TID and plastid-TID, are stably integrated into the Arabidopsis genome we wondered if we would see an enrichment of specific proteins in the CGL20a-TID samples already by TID activity under endogenous levels of biotin. Therefore, we compared proteins between plastid-TID samples and CGL20a-TID samples before the biotin pulse chase (Figure S4). Surprisingly, among the fraction of enriched proteins we only observe proteins which are either cytosolic or facing the cytosol from a different sub-cellular space. Many of them also stay prominently enriched after the biotin pulse chase (Figure S4). Amongst these, the HOP protein group is the strongest enriched before the biotin pulse chase and stays among the three most enriched protein groups after the biotin pulse chase (Figure S4). HSP70/HSP90 organizing proteins (HOP) were reported to associate with plastid pre-proteins in the cytoplasm (Fellerer et al., 2011). This finding prompted us to look in more detail on described core proteins involved in TOC/TIC dependent plastid protein import (Richardson and Schnell, 2019) within our dataset. Indeed, we can observe TOC159 enriched after the pulse chase. Additionally, we find all cytosolic HSP70 protein chaperones (Lin et al., 2001) slightly enriched in the CGL20a-TID samples too (Table S2), indicating that CGL20a-TID preprotein transit to the plastid is assisted by various HSP70s rather than a single isoform. Interestingly we don’t observe any HSP90 proteins to be enriched in the CGL20a-TID dataset except HSP90_6 which harbours a mitochondrial targeting peptide. On the stromal side the only proteins enriched form the plastid import machinery are HSP93/ClpC. This accumulation of plastid protein import machinery components facing the cytosol (Fig S5) leads us to the conclusion that CGL20a-TID seems to be imported more slowly than the plastid-TID control. This finding is surprising since both constructs harbour the same plastid transit peptide. Therefore, we conclude that the peptide sequence of CGL20a itself seems to slow down the plastid import or leads to longer residence in the cytosol. Despite being unexpected, this finding can explain the comparatively vast enrichment of cytoplasmic proteins in the CGL20a-TID samples compared to the plastid-TID samples (FigureS6A).

### The plastid stroma proxiome of CGL20a

In total 79 protein groups were found enriched in the CGL20a-TID samples compared to the control samples after 30 min of Biotin pulse chase (Figure S6A, Table S3). We manually looked for plastid-localized proteins within these 79 protein groups since for many of them only limited consensus accuracy in sub-cellular annotations is retrievable from data provided by global annotation databases such as SUBA5 (Hooper et al., 2017) or the gene ontology resource (GO) (Carbon et al., 2021). We took 3 different data sources into account, 1^st^ sub-cellular localisation annotation through immunological and/or MS/MS analysis of sub-cellular fractionation experiments, 2^nd^ subcellular localisation analysis by analysis of fluorescent protein tagged fusion proteins and/or in-situ hybridisation tags detected either by fluorescence microscopy or electron microscopy, 3^rd^ prediction of a canonical plastid targeting peptide. Only if at least 2 of these sources indicate a plastid localisation we selected the protein as plastid protein, resulting in 19 designated plastid proteins within the initial 79 enriched protein groups (Figure S6a, Table S3). A representation of GO molecular function terms for the 19 plastid proteins shows that ~ 25 % of all assigned GO terms belong to the category “RNA binding” (Figure S6B). In agreement with this, a GO-slim molecular function enrichment analysis within the 19 plastid proteins against the Arabidopsis proteome reports “structural constituent of ribosome” (GO:0003735) and “structural molecular activity” (GO:0005198) as the only enriched GO terms (Table S4). The other proteins have seemingly diverse molecular functions. TOC159 and HSP93/ClpC belong to the plastid protein import machinery. BCCP2 is a component of the plastid localised acetyl-CoA carboxylase complex which was described as one of the first endogenous biotinylation targets in plants (Alban et al., 1994)). CBSX2 and PDE327 are described modulators of photosynthetic electron transport (Yoo et al., 2011 and Vlad et al., 2010). AT2G19940 is a predicted N-acetyl-gamma-glutamyl-phosphate reductase involved in amino acid metabolism. EGY2 is a putative metalloprotease which plays a role in ethylene dependent gravitropism (Chen et al., 2004). AT2G17695 was described as plastid outer envelope protein (Ferro et al., 2010). ACO3 is actually described as an aconitase from mitochondria which was shown to be important for acclimation to submergence stress (Meng et al., 2022). Interestingly, upon submergence an ACO3-YFP fusion protein is detected also inside plastids (Meng et al., 2022). FBA1 is described as plastidial fructose-1,6-bisphosphate aldolase (Carrera et al., 2021) and EMB3119 is a putative ribose 5-phosphate isomerase displaying an embryo-lethal knock out phenotype (Meinke, 2019). Interestingly, EMB3119 has also been described as thioredoxin interacting protein (Marchand et al., 2006), which would provide an interesting link to CBSX2 in redox regulation (Yoo et al., 2011). GUN2 encodes a heme-oxygenase (Wang et al., 2022) necessary for coupling the expression of certain nuclear genes to the functional state of the chloroplast (Mochizuki et al., 2001). The six remaining proteins belong all to the RNA binding protein family.

### CGL20a resides in a nexus of RNA binding proteins

Reiter et al. reported that a fraction of CGL20a protein co-migrates with high molecular weight ribosome containing fractions in a sucrose density gradient (Reiter et al., 2020). Since we found the go category “structural constituent of the ribosome” enriched in our data we decided to follow all plastid ribosomal proteins in our data, irrespective of the arbitrarily chosen 5 % FDR rate. First, we obtained a list of described plastid ribosomal proteins from various manuscripts (Tiller and Bock, 2014; Bieri et al., 2016; Graf et al., 2016; Ahmed et al., 2017;) and followed them in our enrichment data (Figure 5A). From a total of 53 plastid ribosomal proteins, we identified 41 in our dataset. We observe that during the 30 min biotin pulse chase the majority of the plastid ribosomal proteins seems to move to the left side in the volcano blot, indicating CGL20a-TID dependent enrichment (Figure 5A). These data suggest that CGL20a resides somewhere in the vicinity of the plastid ribosome. Reiter et al. also observed defects in the hidden break formation of the 23S rRNA (Leaver, 1973) in *cgl20a/b* double knock out mutants (Reiter et al., 2020). The Deadbox RNA Helicase RH39 is described to play a key role in the hidden break formation (Nishimura et al., 2010). We detected RH39 in our data set, however, it is clearly less strongly enriched in a CGL20a-TID dependent way than most of the plastid ribosomal proteins. Therefore, it seems to be unlikely that CGL20a is directly involved in hidden break formation but rather seems to exert influence on the process indirectly. Besides the genuine plastid ribosomal proteins, we observed two more RNA interacting proteins related to ribosomal function, RUG2 and a SPOUT methyltransferase. RUG2 belongs to the family of metazoan mitochondrial transcription termination factors (mTERFs) with various molecular functions (Robles and Quesada, 2021). In Arabidopsis RUG2 is observed to dually localise to chloroplasts and mitochondria. A *rug2* functional knock out was described as severely perturbed in mitochondria and chloroplast development and leading to a strongly increased transcription of nuclear encoded plastid RNA polymerase (Quesada et al., 2011). In parallel, splicing deficiencies for plastid transcribed rps12, rpl2, atpF and the clpP mRNAs were described in *rug2* mutants along with its inability to accumulate mature ribosomal RNA (Babiychuk et al., 2011). SPOUT methyltransferases are described to exert several functions. Members of this family were reported to methylate tRNAs and rRNAs, and in exceptional cases also proteins thereby mainly influencing transcriptional efficiency (Strassler et al., 2022). Unfortunately, no exact molecular mechanisms on the function of these two proteins are yet described but our data for the first time places these two proteins in the vicinity of the plant plastid ribosome, thereby, building the basis for further investigation.

**Figure 5.**
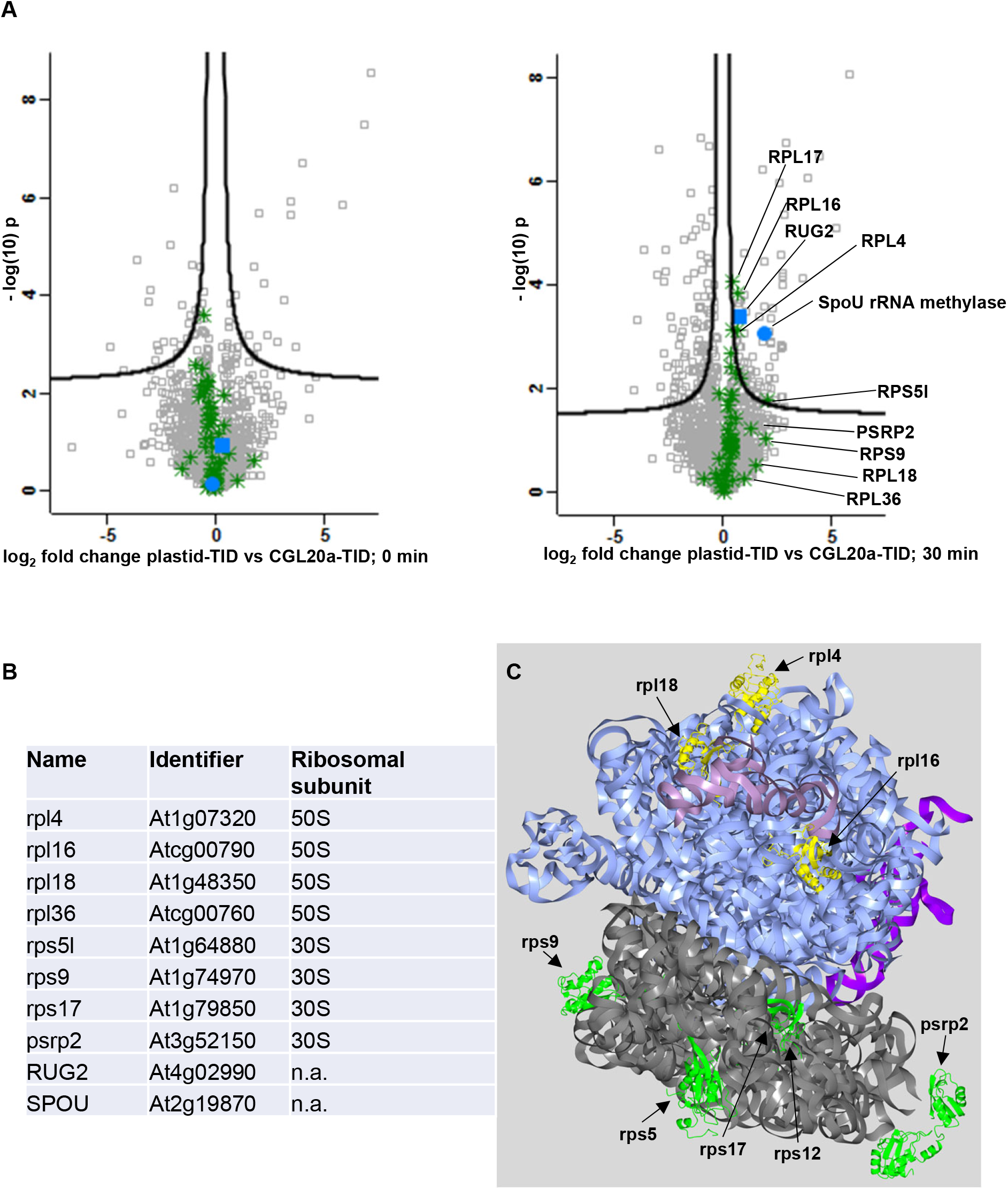
CGL20a-TID resides in a nexus of RNA binding proteins. **A**, Left panel, volcano plot of SA enriched protein groups from *plastid-TID* vs *CGL20a-TID* samples after 0 min and after 30 min (right panel) of biotin treatment. All green stars mark plastid ribosomal proteins. The blue circle marks SpoU type rRNA methyltransferase and the blue square marks RUG2. Solid black lines indicate significance thresholds for an FDR of 0.05. 5 biological replicates were analysed for each genotype, condition and timepoint. **B**, list of the 10 most enriched plastid RNA bidning proteins in *GL20a-TID* samples through out the 30 min biotin pulse chase. **C**, a rendered model of the plastid ribosome according to (Achmed et al., 2017) blue, 23SrRNA; pink, 5SrRNA; violet, 4.5SrRNA; grey, 16SrRNA; .green, proteins from B associated to the small subunit; yellow, proteins from B associated to the large subunit.

## Discussion

### TurboID based proximity labelling inside plastids of plants and algae works

Despite successful application of TID-based proximity labelling in plant cells at different sub cellular localisations (Mair et al., 2019; Tang et al., 2022; Arora et al., 2020; Xu et al 2021), it has always been questioned if it would work in plastids of higher plant cells as well. Reports on the free biotin concentration suggested only marginal amounts inside chloroplasts (Badelt et al 1993). This might have been a reason why biotin ligase-based proximity labelling in plastids has not been approached yet. Here we present data that application of 50 μM exogenous biotin leads to measurable increase of plastid targeted TID activity after 30 minutes (Figure 3). Under these conditions we observe that plastid targeted TID mediates the enrichment of the majority of proteins after SA pulldowns compared to a Col0 control (Figure 4 and Figure S3). Done in parallel, two independent studies report functional TID-dependent proximity labelling in plastids of *Chlamydomonas reinhardtii* (Lau et al., submitted; Kreis et al., submitted). Together, these data suggest that TID-based proximity labelling works in plastids of diverse species. At the same time, we also observe important differences between these species. Whereas for Arabidopsis plastids we find addition of 50 μM exogenous biotin enough to obtain sufficient TID dependently enriched proteins for analysis, both studies in Chlamydomonas report the need of supplementing exogenous biotin between 0.5 and 1 mM for efficient labelling inside plastid stroma (Kreis et al., submitted). In case of pyrenoid localised TID labelling even concentrations > 1mM were found to be necessary (Lau et al., submitted).

In their manuscripts, Kreis and colleagues as well as Lau and colleagues observe that high expression of TID leads to alterations in the endogenous biotinylation pattern. This effect was attributed to competition between TID and the endogenous biotin ligases for endogenous biotin as it could be largely reverted by addition of exogenous biotin (Kreis et al., submitted; Lau et al., submitted). In Arabidopsis we were not able to see such an obvious effect, and both *cgl20a-TID* and *plastid-TID* lines did not show any obvious phenotype compared to Col0 after 4 generations of propagation indicating that constitutive presence of TID, at least at moderate levels, does not severely interfere with endogenously required biotinylation.

### The CGL20a-TID proxiome

The previous study by Reiter et al. (2020) suggested that the strong virescent phenotype of *cgl20a/b* mutants is mostly based on impaired ribosome processing via a defect in hidden break formation. How hidden break formation in plastid ribosomes occurs is currently not completely elucidated. Interestingly, also for *rug2* mutants a defect in maturation of plastid ribosomal RNA has been reported (Babiychuk et al., 2011). Also, the RNA helicase RH39 was shown to take part in this process (Nishimura et al., 2010). However, our data does not show CGL20a-dependent enrichment of RH39. Additionally, RUG2 very specifically localises to plastid nucleoid structures (Babiychuk et al., 2011) whereas CGL20a seems to be dispersed in the plastid stroma (Figure 1C). Taking all this data together suggests that plastid rRNA maturation is an elaborate process taking place in several steps at different sites in the plastid stroma. Our observation, that plastid ribosomal proteins associated to the large and small subunit, were evenly enriched in our data-set is in agreement with previous data (Reiter et al., 2020), suggesting that CGL20a acts at the late stages of ribosome maturation.

Chloroplast genomes encode ~37 proteins that integrate into the thylakoid membrane and already about half a century ago it was reported that chloroplast ribosomes were bound to the unstacked regions of the thylakoid membranes in Chlamydomonas (Chua et al., 1973; Margulies and Michaels, 1974), which was later also found for higher plants (Yamamoto et al., 1981). More recently, a genome-wide, high-resolution analysis of the partitioning of chloroplast ribosomes between membrane and soluble fractions in maize seedlings revealed that approximately half of the chloroplast-encoded thylakoid proteins integrate cotranslationally and half integrate posttranslationally (Zoschke and Barkan, 2015). Translation initiates off the thylakoid membrane and ribosomes synthesizing a subset of membrane proteins subsequently become attached to the membrane in a nuclease-resistant fashion. Several central photosynthetic proteins, such as the PSII PsbA (D1) subunit in Arabidopsis, are cotranslationally inserted into the thylakoid membrane mediated by the chloroplast signal recognition particle subunit cpSRP54 (Hristou et al., 2019). In this context, the finding that PsbA and other PSII subunits are severely affected at protein level in *cgl20a* mutants (Reiter et al., 2020; Leonardelli, 2017), and the CGL20a-dependent enrichment of thioredoxin interacting proteins CBSX2 and EMB3119 might be an indication that CGL20a is associated to ribosomes associated to the thylakoid membrane.

### The future of TurboID based proximity labelling in plastids

One of our main goals was to show that TID based proximity labelling can work with comparatively minute amounts of plant material (12 seedlings per sample, 500 μg of total protein). This now allows to probe for protein proxiomes in plastids within highly dynamic samples such as early developing seedlings during chloroplast development of meristematic tissues where the low amount of sample material prevented large-scale protein interaction analysis so far. Mair and colleagues elegantly showed that by using cell line specific promoters for bait-TID constructs populations of FAMA transcription factor vicinal proteins are determinable within the guard cell progenitor lineage (Mair et al., 2019) *in vivo* in the context of the whole plant. It is now well established, that plastids are highly dynamic organelles, able to differentiate into numerous specialised forms within a tissue dependent context (Liebers et al., 2022). By expressing plastid targeted TID constructs under control of cell line-specific promoters it will be possible to follow bait proxiome changes specifically inside plastids as they are embedded in their endogenous plant context. All without the need of separating different cell populations by protoplasting and subsequent cell sorting or comparatively laborious tissue specific sampling by laser micro dissection. In addition, labelling times of 30 minutes should also allow to follow developmental transitions of plastids such as differentiation of chloroplasts, sensory plastids, amyloplasts, eoplasts etc. (Liebers et al., 2017). Still, especially for plants it would be very important to improve labelling speed of promiscuous biotin ligases. Arora and colleagues showed that robustness of results with TID increases significantly at labelling times of hours rather than minutes at temperatures above 22 °C (Arora et al., 2020). To our knowledge, there is also no report on the activity of promiscuous biotin ligases in plants at temperatures below 20 °C. Given the fact that plants in temperate climate zones have to deal with temperature changes from below zero to more than 30 °C throughout the year it would be valuable to have a proximity labelling tool similar to TID which works at sufficient speed between 4 and 20 °C too.

## Materials and Methods

### Molecular cloning

DNA constructs used in this study were assembled via PCR based cloning with NEB builder. All primers used in this study were automatically designed by the NEB builder software (https://nebuilder.neb.com) from the company New England Biolabs. For assembling cgl20a-TID the cgl20a genomic locus including 2kb immediately upstream of the 5’UTR was PCR amplified from Arabidopsis thaliana Col0 genomic DNA. The cgl20a STOP codon in Exon2 was replaced by the TID-YFP cassette from the vector pDONR-P2R-P3_R2-Turbo-mVenus-STOP-L3_(#3409) kindly provided by Andrea Mair (Mair et al., 2019). Both PCR fragments were assembled together with a pBIB_Hyg plant transformation vector backbone resulting in the plasmid cgl20a-TID_pBIB_Hyg (supplementary data1) carrying a Hygromycin resistance cassette for selection in plants. To obtain the plastid-TID control construct the promoter, 5’UTR and Exon1 including the plastid targeting peptide was PCR amplified from A.thaliana Col0 genomic DNA. The ATG encoding methionine immediately after the cgl20a plastid targeting peptide was replaced by the same TID-YFP cassette as described above. Both PCR fragments were assembled together with the pBIB Hyg backbone to result in a plant transformation vector targeting TID-YFP to the plastid stroma by the use of the cgl20a plastid targeting peptide (supplementary data 2).

### Plant transformation

Both vectors were transformed into Agrobacteria strain GV3101. By agro bacterium mediated floral dip transformation (Clough and Bent, 1998) the cgl20a-TID construct was transformed in to a *cgl20a/b* functional double knock out (dko) in *Col0* background. The plastid-TID control construct was transformed into Col0. Initial F1 transformants were screened on agar plates (4.4 g/l Murashige and Skoog salts, 1 % (w/v) sucrose, 0.7 % (w/v) plant agar, pH 5.8) containing Hygromycin (15 mg/l), Vancomycin (100 mg/l) and Cefatoxin (100 mg/l). Both Vancomycin and Cefatoxin where only used in the F1 screen to avoid accidental bacterial growth. For F1-3 screens seeds were stratified on plates in dark at 4°C for 4 days. Subsequently plates were put into light under long day conditions (16h light/ 8 hours dark) at 22 °C. Hygromycin resistant plants were selected after 12 days and transferred to soil for seed propagation. In case of CGL20a-TID we also selected for complementation of the *cgl20a/b* dko phenotype to finally obtain plants similar in greening speed to *Col0*. For all experiments, plants of the F4 generation were used.

### Plant growth and Biotin pulse chase

Surface sterilized seeds were stratified on MS plates supplemented with 1 % (w/v) sucrose for 4 days and then grown for 7 days under long day conditions. For the biotin pulse chase 12 seven-day old seedlings, for each condition and genotype, were transferred in to 3 ml liquid 1/2 MS medium (pH 6.5) in a 6 well plate and left shaking for 1 hour. Immediately after this hour the 0 min timepoint control samples were washed 3 times for 2 minutes with ice cold water. After the last wash step seedlings were gently surface dried between tissue and frozen in liquid nitrogen. For the biotin pulse chase the MS medium was exchanged for MS medium supplemented with 50 μM of Biotin. After the indicated time points seedlings were washed, dried and frozen as described above.

### Protein extracts for immuno blots

Frozen plant material was homogenised by shaking the seedlings with 2 metalbeads, 2.5 mm in diameter per 2 ml microreaction sample tube in a tissue lyser II (Quiagen) at a frequency of 28 Hz for 1 minute. The obtained powder was thawed and resuspended in SDS-Page loading buffer (50 mM Tris/Cl pH 6.8, 2 % (w/v) SDS, 10 % (v/v glycerol, 2.5 % (v/v) 2-mercaptoethanol, 0.025 % (w/v) bromophenol blue). Reduction of samples was aided by incubation at 95 °C for 5 minutes. Subsequently samples were separated by SDS PAGE and blotted onto immobilon-P PVDF membaren (Millipore) by using a Trans-Blot Turbo transfer system from BioRad according to the manufacturer’s instructions. Membranes were blocked in 1 % (w/v) BSA in TBS-Tween (0.05 % v/v) for 1 hour. Exposure to primary antibodies (rat - anti GFP 3H9-Proteintech or mouse – anti Biotin B7653-Sigma-Aldrich) was done for 2 hours at 25°C. Exposure to secondary antibodies (anti rat-HRP, Cytiva #NA935 or anti mouse-HRP, Cytiva #NA931) was done for 1 hour at 25°C. Western blots were developed by the use of western bright Sirius chemiluminescent detection kit ADVANSTA #K12043 and BioRad ChemiDoc imaging system.

### Streptavidin mediated enrichment of biotinylated proteins, digest and peptide cleanup

All subsequent manipulation steps were done in lo-bind microreaction tubes (#EP0030108116, Merck) to avoid protein/peptide loss by unspecific adherence to the tube wall. Frozen, homogenised plant material, as described above, was thawed and resuspended in 500 μl extraction buffer (50 mM Tris/Cl pH 7.5; 150 mM NaCl, 0.5 mM EDTA; 0.5 % NP40 substitute; 1 mM DTT; 1 Tab/25 ml protease inhibitor complete tablette (#11836170001 Roche)). The resulting suspension was incubated 5 min on ice and then centrifuged 2 min, 20.000 g, 4 °C. Protein concentrations of the soluble proteins in the supernatant were determined by BCA assay (Pierce Microplate BCA Protein assay kit, # 23252) and adjusted to 1 mg/ml. 500 μl of that total protein (500 μg) extract were subjected to gel filtration via PD MiniTrap G-25 (#28918007, Cytiva) columns equilibrated in extraction buffer to remove free endogenous biotin interfering with the subsequent streptavidin (SA) biotinylated-protein enrichment.

20 μl of in extraction buffer equilibrated streptavidin magnetic beads (S1420, NEB) were added to 1 ml of the free biotin depleted protein extracts. The suspension was incubated for 60 min at 4°C and constant mixing on a rotator. Subsequently, the beads were washed four times with washbuffer (20 mM Tris/Cl pH7.5; 500 mM NaCl, 0.5 mM EDTA) and transferred to a fresh microreaction tube.

On beads digests of proteins and peptide clean-up for mass spectrometry (MS) was done with an iST proteomics preparation kit (P.O.00027, Preomics) according to the manufacturer’s instructions. The adjustable parameters in brief, 50 μl of “Lyse” solution was added to each sample of beads and incubated at 60 °C and shaking at 1400 rpm for 10 min. 50 μl of protease containing “Digest” solution were added to the beads and incubated at 37 °C, 2 hours under constant shaking at 1400 rpm. Digest was stopped by adding 100 μl of “STOP” solution mixed and applied to the peptide clean-up cartridge. Peptides were bound to the matrix by centrifugation of the cartridge at 3800 g, 3 min, 25 °C. Peptides were sequentially washed with 200 μl “Wash1” and “Wash2” at the same centrifugation set up. Peptides were sequentially eluted in 2 times 100 μl “Elute” solution, collected in a 1.5 ml microreaction tube. Obtained peptides were dried in a speed vac system (Concentrator plus, Eppendorf) at 45 °C until the solvent completely evaporated (~ 2.5 hours).

### LC-MS analysis

#### Nano-liquid choromatography (LC)

Purified tryptic peptides were dissolved in 0.1% formic acid (FA) solution (v/v) (in high purity water (MilliQ)). Then 1μg of peptides was separated by an online reversed-phase (RP) HPLC (Thermo Scientific Dionex Ultimate 3000 RSLC nano LC system) connected to a benchtop Quadrupole Orbitrap (Q-Exactive Plus) (QE Plus) mass spectrometer (Thermo Fisher Scientific). The online separation was performed on an Easy-Spray analytical column (PepMap RSLC C18, 2 μm, 100 Å, 75 μm i.d. × 50 cm, Thermo Fisher Scientific) with an integrated emitter. The Column was heated at 55°C. The flow rate was set to 300 μL/min. The LC gradient was a two-hour gradient method and was set to 5 - 50% buffer B (v/v) [79.9% ACN, 0.1% formic acid (FA), 20% Ultra high purity (MilliQ)] for 110 minutes and then to 80% buffer B over 5 minutes. The buffer A (v/v) was 0.1% FA in high purity water (MilliQ).

### MS

Benchtop-Quadrupole Orbitrap (Q-Exactive Plus)

LC eluent was introduced into the mass spectrometer through an Easy-Spray ion source (Thermo Scientific). The emitter was operated at 1.9 kV. The mass spectra were measured in positive ion mode applying a top fifteen data-dependent acquisition (DDA). A full mass spectrum was set to 70,000 resolution at m/z 200 [AGC target at 1e6, maximum injection time (IT) of 120 ms and a scan range 400-1600 (m/z)]. The MS scan was followed by a MS/MS scan at 17,500 resolution at m/z 200 [Automatic Gain Control (AGC) target at 5e4, 1.6 m/z isolation window and maximum IT of 80 ms]. For MS/MS fragmentation, normalized collision energy (NCE) for higher energy collisional dissociation (HCD) was set to 27%. Dynamic exclusion was at 40 s. Unassigned and +1, +7, +8 and > +8 charged precursors were excluded. The intensity threshold was set to 6.3e3. The isotopes were excluded.

### MS data analysis (LFQ – label free quantification)

Protein identification and LFQ were done with Maxquant (v2.1.4.0) (Tyanova et al., 2016a). Default settings with minor modifications were used for the peptide identification and LFQ in Maxquant. Since the background proteomes and TID labelling efficiency were observed to be similar by western blot analysis (Fig 3) all data obtained from samples at different time points and genotypes (0 min, 30 min, Col0, plastid-TID and cgl20a-TID) were analysed in parallel in Maxquant. Within the group specific parameters, the “oxidation (M)” and “acetylation (Protein N-term)” were selected as variable modifications and “carbamidomethyl (C)” as fixed modification. The maximum number of modifications was set to 5. Digestion mode was set to “specific” and enzymes “Trypsin/P” and “LysC/P” with 2 max. missed cleavage sites were selected. For LFQ the normalisation type “classic” was selected and Fast LFQ was ticked setting min. number of neighbours to “3” and max. number of neighbours to “6”. As search template the araport 11 Arabidopsis protein database (v20220103, accessed via TAIR www.arabidopsis.org) including the polypeptide sequences of the respective TID constructs was used in parallel the internal maxquant protein contaminations list was used to identify frequent protein contaminants in shotgun proteomics samples. For Peptide quantification “Unique + razor” was selected and only unmodified and “oxidation (M)”, “acetyl (Protein N-term)”, “carbamidomethyl (C)” modified peptides were selected plus discard unmodified counterpart peptides was set to “yes”. For identification the FDR was set to 0.01. All other options were left default.

Explorative analysis of Maxquant LFQ data output “proteinGroups.txt” was performed in the Perseus software package (v2.0.3.0) (Tyanova et al., 2016b). First results were filtered for the following three categories, “proteins only identified by site”, “reverse” and “potential contaminant”. Replicate samples were grouped according to the biological condition they were obtained from. All remaining values were log 2 transformed, only protein groups identified in at least 1 one sample group with at least three valid values out of five replicates were retained for further analysis. In this subset missing values were imputed with Perseu’s default “Replace missing values from normal distribution function” (width = 0.3, down shift = 1.8, mode = total matrix) for further statistical analysis. Volcano plot representations of two-sided T-tests were created in Preseus, exported as image and additional labelling, of individual identifiers was done in MS power point (Figures 5 and Figures S4). For orientation the arbitrarily chosen 5 % FDR “cut off” – line is depicted in the volcano blots.

### GO term analysis

The functional Categorization by annotation for GO molecular function (Figure S6B) was obtained by querying the 19 CGL20a-TID dependently enriched proteins in the TAIR (www.arabidopsis.org) in the bulk data GO-term retrieval tab.

GO molecular function enrichment analysis was performed online at the Gene Ontology resource (http://geneontology.org/) (Carbon et al., 2021) by querying the CGL20a-TID dependently enriched proteins against the TAIR10 proteomic database.

### Rendering of plastid ribosomal structure

The structure of the cryo-EM derived plastid ribosome, PDBID: 5×8P (www.rcsb.org) (Berman et al., 2000), deposited by Ahmed and co-workers (Ahmed et al., 2017) was used as template to render Figure 5C with the software package CCP4MG (www.ccp4.ac.uk) (McNicholas et al., 2011).

### Imaging

Images of seedlings grown on plates (Figure 1A) were recorded with a Canon EOS D90 camera system equipped with an EPS 17 – 25 mm wide field lens.

Fluorescence images (Figure 1C) were recorded on a Leica SP8 system. Immediately before microscopy seedlings were infiltrated with H2O to fill the air spaces of the parenchym thereby, creating a more homogenous refractive index across the sample. Fluorophores were excited with a RYB-Argon laser at 514 nm. YFP signal was recorded between 525 and 540 nm with a HyD detector. Plastid autofluorescence was recorded in parallel with a conventional photomultiplier from 680 to 730 nm.

### Transmission electron microscopy

For ultrastructural analysis 2 mm^2^ sections of at least three leaves from mature rosette leaves of WT and *cgl20a/b* dko plants were used for combined conventional and microwave-proceeded fixation (paraformaldehyde, glutaraldehyde and osmium tetroxide), (details see in supplementary methods 1). The procedures for dehydration, resin embedding, ultrathin sectioning, and ultrastructural analysis have been previously described (Kraner et al., 2017).

## Supplementary data list

Supplementary figure S1 – segregation analysis summary

Supplementary figure S2 – Western blots showing expression of proximity-labelling constructs

Supplementary figure S3 – protein enrichment after biotin pulse-chase in Col0

Supplementary figure S4 – CGL20a-specifically enriched protein groups before pulse-chase

Supplementary figure S5 – scheme of core components of the plastid import machinery

Supplementary figure S6 – CGL20a-dependent enriched plastid proteins

Supplementary data set 1 – protein groups identified in all samples (basis for enrichment analysis)

Supplementary data set 2 – table with plastid import factors and their CGL20a dependent enrichment

Supplementary data set 3 – Table of CGL20a dependent enriched proteins (plastid proteins are highlighted in green)

Supplementary data set 4 – Enrichment analysis GO_slim_molecular function

Supplementary data set 5 – Vector map of CGL20-TID plant transformation vector

Supplementary data set 6 – Vector map of plastid-TID plant transformation vector

Supplementary data set 7 – Table listing details for fixation of samples for transmission electron microscopy

## Acknowledgements

We thank the Cell Imaging & Utrastructure Research facility (CUIS) of the Faculty of Life Sciences at the University of Vienna for providing access to fluorescence microscopes and Andreas Bachmair (Max Perutz Labs, University of Vienna) for sharing lab space. We acknowledge funding by the EU Horizon 2020 research and innovation project ADAPT (GA 2020 862-858).

## Author Contributions

Bernhard Wurzinger designed and performed the research; analyzed data and wrote the paper; Simon Stael performed research and provided tools; Manuela Leonardelli performed research and provided tools; Carlo Perolo performed research; Michael Melzer performed research; Palak Chaturvedi performed research, Leila Afjehi-Sadat performed research; Wolfram Weckwerth contributed tools; Markus Teige designed research, analysed data and wrote the paper.

## References

Alban, C., Baldet, P., and Douce, R. (1994). No TitlLocalization and characterization of 2 structurally different forms of acetyl-CoA carboxylase in young pea leaves, of which one is sensitive to aryloxyphenoxypropionate herbicidese. Biochem. J. 300: 557–565.

Alban, C. (2011). Biotin (Vitamin B8) Synthesis in Plants. Adv. Bot. Res. 59: 39–66.

Ahmed, T., Shi, J., and Bhushan, S. (2017). Unique localization of the plastid-specific ribosomal proteins in the chloroplast ribosome small subunit provides mechanistic insights into the chloroplastic translation. Nucleic Acids Res. 45: 8581–8595.

Arora, D. et al. (2020). Establishment of proximity-dependent biotinylation approaches in different plant model systems. Plant Cell 32: 3388–3407.

Babiychuk, E., Vandepoele, K., Wissing, J., Garcia-Diaz, M., De Rycke, R., Akbari, H., Joubès, J., Beeckman, T., Jänsch, L., Frentzen, M., Van Montagu, M.C.E., and Kushnir, S. (2011). Plastid gene expression and plant development require a plastidic protein of the mitochondrial transcription termination factor family. Proc. Natl. Acad. Sci. U. S. A. 108: 6674–6679.

Baldet, P., Alban, C., Axiotis, S., and Douce, R. (1993). Localization of Free and Bound Biotin in Cells from Green Pea Leaves. Arch. Biochem. Biophys. 303: 67–73.

Berman, H.M., Westbrook, J., Feng, Z., Gililand, G., Bhat, T.N., Weissig, H., Shindyalov, I.N., and Bourne, P.E. (2000). The Protein Data Bank. Nucleic Acids Res. 28: 235–242.

Bieri, P., Leibundgut, M., Saurer, M., Boehringer, D., and Ban, N. (2017). The complete structure of the chloroplast 70S ribosome in complex with translation factor pY. EMBO J. 36: 475–486.

Cackett, L., Luginbuehl, L.H., Schreier, T.B., Lopez-Juez, E., and Hibberd, J.M. (2022). Chloroplast development in green plant tissues: the interplay between light, hormone, and transcriptional regulation. New Phytol. 233: 2000–2016.

Carbon, S. et al. (2021). The Gene Ontology resource: Enriching a GOld mine. Nucleic Acids Res. 49: D325–D334.

Carrera, D.Á., George, G.M., Fischer-Stettler, M., Galbier, F., Eicke, S., Truernit, E., Streb, S., and Zeeman, S.C. (2021). Distinct plastid fructose bisphosphate aldolases function in photosynthetic and non-photosynthetic metabolism in Arabidopsis. J. Exp. Bot. 72: 3739–3755.

Charuvi, D., Kiss, V., Nevo, R., Shimoni, E., Adam, Z., and Reich, Z. (2012). Gain and loss of photosynthetic membranes during plastid differentiation in the shoot apex of arabidopsis. Plant Cell 24: 1143–1157.

Chen, G., Bi, Y.R., and Li, N. (2005). EGY1 encodes a membrane-associated and ATP-independent metalloprotease that is required for chloroplast development. Plant J. 41: 364–375.

Chua N.H., Blobel G., Siekevitz P., Palade G.E. (1973). Attachment of chloroplast polysomes to thylakoid membranes in Chlamydomonas reinhardtii. Proc Natl Acad Sci USA. 70: 1554–1558.

Clough, S.J. and Bent, A.F. (1998). Floral dip: A simplified method for Agrobacterium-mediated transformation of Arabidopsis thaliana. Plant J. 16: 735–743.

Dubreuil, C., Jin, X., Barajas-López, J. de D., Hewitt, T.C., Tanz, S.K., Dobrenel, T., Schröder, W.P., Hanson, J., Pesquet, E., Grönlund, A., Small, I., and Strand, Å. (2018). Establishment of photosynthesis through chloroplast development is controlled by two distinct regulatory phases. Plant Physiol. 176: 1199–1214.

Fellerer, C., Schweiger, R., Schöngruber, K., Soll, J., and Schwenkert, S. (2011). Cytosolic HSP90 cochaperones HOP and FKBP interact with freshly synthesized chloroplast preproteins of arabidopsis. Mol. Plant 4: 1133–1145.

Ferro, M. et al. (2010). AT-CHLORO, a comprehensive chloroplast proteome database with subplastidial localization and curated information on envelope proteins. Mol. Cell. Proteomics 9: 1063–1084.

Graf, M., Arenz, S., Huter, P., Dönhöfer, A., Nováček, J., and Wilson, D.N. (2017). Cryo-EM structure of the spinach chloroplast ribosome reveals the location of plastid-specific ribosomal proteins and extensions. Nucleic Acids Res. 45: 2887–2896.

Hooper, C.M., Castleden, I.R., Tanz, S.K., Aryamanesh, N., and Millar, A.H. (2017). SUBA4: The interactive data analysis centre for Arabidopsis subcellular protein locations. Nucleic Acids Res. 45: D1064–D1074.

Hristou A., Gerlach I., Stolle D.S., Neumann J., Bischoff A., Dünschede B., Nowaczyk M.M., Zoschke R., Schünemann D. (2019). Ribosome-Associated Chloroplast SRP54 Enables Efficient Cotranslational Membrane Insertion of Key Photosynthetic Proteins. Plant Cell 31: 2734–2750.

Kraner M., Link K., Melzer M., Ekici A.B., Uebe S., Tarazona Corrales P., Feussner I. & Sonnewald U. (2017) Choline transporter-like1 (CHER1) is crucial for plasmodesmata maturation in Arabidopsis thaliana. Plant J. 89: 394–406

Kreis, E., Koenig, K., Sommer, F., and Schoroda M. TurboID reveals the interaction networks of CGE1, VIPP1, and VIPP2 in Chlamydomonas reinhardtii. submitted

Lau, C.S., Dowle, A., Thomas, G., Girr, P., and Mackinder, L.C. The phase separated CO2-fixing pyrenoid proteome determined by TurboID. submitted

Leaver, C.J. (1973). Molecular integrity of chloroplast ribosomal ribonucleic acid. Biochem. J. 135: 237–240.

Leonardelli, M. (2017). Characterization of two novel chloroplast proteins with a potential role in chloroplast development and function. PhD thesis University of Vienna, 2017.

Liebers, M., Grübler, B., Chevalier, F., Lerbs-Mache, S., Merendino, L., Blanvillain, R., and Pfannschmidt, T. (2017). Regulatory shifts in plastid transcription play a key role in morphological conversions of plastids during plant development. Front. Plant Sci. 8: 1–8.

Liebers, M., Cozzi, C., Uecker, F., Chambon, L., Blanvillain, R., and Pfannschmidt, T. (2022). Biogenic signals from plastids and their role in chloroplast development. J. Exp. Bot. 73: 7105–7125.

Lin, B.L., Wang, J.S., Liu, H.C., Chen, R.W., Meyer, Y., Barakat, A., and Delseny, M. (2001). Genomic analysis of the Hsp70 superfamily in Arabidopsis thaliana. Cell Stress Chaperones 6: 201–208.

Mair, A., Xu, S.L., Branon, T.C., Ting, A.Y., and Bergmann, D.C. (2019). Proximity labeling of protein complexes and cell type specific organellar proteomes in Arabidopsis enabled by TurboID. Elife 8: 1–45.

Mair, A, Bergmann, D. (2022) Advances in enzyme-mediated proximity labeling and its potential for plant research. Plant Physiol. 188: 756–768.

Marchand, C., Le Maréchal, P., Meyer, Y., and Decottignies, P. (2006). Comparative proteomic approaches for the isolation of proteins interacting with thioredoxin. Proteomics 6: 6528–6537.

Margulies M.M., Michaels A. (1974). Ribosomes bound to chloroplast membranes in Chlamydomonas reinhardtii. J Cell Biol. 60: 65–77.

McNicholas, S., Potterton, E., Wilson, K.S., and Noble, M.E.M. (2011). Presenting your structures: The CCP4mg molecular-graphics software. Acta Crystallogr. Sect. D Biol. Crystallogr. 67: 386–394.

Meinke, D.W. (2020). Genome-wide identification of EMBRYO-DEFECTIVE (EMB) genes required for growth and development in Arabidopsis. New Phytol. 226: 306–325.

Meng, X., Li, L., Pascual, J., Rahikainen, M., Yi, C., Jost, R., He, C., Fournier-Level, A., Borevitz, J., Kangasjärvi, S., Whelan, J., and Berkowitz, O. (2022). GWAS on multiple traits identifies mitochondrial ACONITASE3 as important for acclimation to submergence stress. Plant Physiol. 188: 2039–2058.

Mochizuki, N., Brusslan, J.A., Larkin, R., Nagatani, A., and Chory, J. (2001). Arabidopsis genomes uncoupled 5 (GUN5) mutant reveals the involvement of MG-chelatase H subunit in plastid-to-nucleus signal transduction. Proc. Natl. Acad. Sci. U. S. A. 98: 2053–2058.

Nishimura, K., Ashida, H., Ogawa, T., and Yokota, A. (2010). A DEAD box protein is required for formation of a hidden break in Arabidopsis chloroplast 23S rRNA. Plant J. 63: 766–777.

Pipitone, R., Eicke, S., Pfister, B., Glauser, G., Falconet, D., Uwizeye, C., Pralon, T., Zeeman, S.C., Kessler, F., and Demarsy, E. (2021). A multifaceted analysis reveals two distinct phases of chloroplast biogenesis during de-etiolation in arabidopsis. Elife 10: 1–32.

Quesada, V., Sarmiento-Mañús, R., González-Bayón, R., Hricová, A., Pérez-Marcos, R., Graciá-Martínez, E., Medina-Ruiz, L., Leyva-Díaz, E., Ponce, M.R., and Micol, J.L. (2011). Arabidopsis RUGOSA2 encodes an mTERF family member required for mitochondrion, chloroplast and leaf development. Plant J. 68: 738–753.

Reiter, B., Vamvaka, E., Marino, G., Kleine, T., Jahns, P., Bolle, C., Leister, D., and Rühle, T. (2020). The Arabidopsis protein CGL20 is required for plastid 50S ribosome biogenesis. Plant Physiol. 182: 1222–1238.

Richardson, L.G.L. and Schnell, D.J. (2020). Origins, function, and regulation of the TOC-TIC general protein import machinery of plastids. J. Exp. Bot. 71: 1226–1238.

Robles, P. and Quesada, V. (2021). Research progress in the molecular functions of plant mTERF proteins. Cells 10: 1–16.

Strassler, S.E., Bowles, I.E., Dey, D., Jackman, J.E., and Conn, G.L. (2022). Tied up in knots: Untangling substrate recognition by the SPOUT methyltransferases. J. Biol. Chem. 298: 102393.

Tadini, L., Jeran, N., Peracchio, C., Masiero, S., Colombo, M., and Pesaresi, P. (2020). The plastid transcription machinery and its coordination with the expression of nuclear genome: Plastid-encoded polymerase, nuclear-encoded polymerase and the genomes uncoupled 1-mediated retrograde communication. Philos. Trans. R. Soc. B Biol. Sci. 375.

Tang, Y., Ho, M.I., Kang, B.-H., and Gu, Y. (2022). GBPL3 localizes to the nuclear pore complex and functionally connects the nuclear basket with the nucleoskeleton in plants. PLOS Biol. 20: e3001831.

Tiller, N. and Bock, R. (2014). The translational apparatus of plastids and its role in plant development. Mol. Plant 7: 1105–1120.

Tissot, G., Douce, R., and Alban, C. (1997). Evidence for multiple forms of biotin holocarboxylase synthetase in pea (Pisum sativum) and in Arabidopsis thaliana: subcellular fractionation studies and isolation of a cDNA clone. Biochem. J. 323: 179–188.

Tyanova, S., Temu, T., and Cox, J. (2016a). The MaxQuant computational platform for mass spectrometry-based shotgun proteomics. Nat. Protoc. 11: 2301–2319.

Tyanova, S., Temu, T., Sinitcyn, P., Carlson, A., Hein, M.Y., Geiger, T., Mann, M., and Cox, J. (2016b). The Perseus computational platform for comprehensive analysis of (prote)omics data. Nat. Methods 13: 731–740.

Vlad, D., Rappaport, F., Simon, M., and Loudet, O. (2010). Gene transposition causing natural variation for growth in Arabidopsis thaliana. PLoS Genet. 6: 21.

Wang, J., Li, X., Chang, J.W., Ye, T., Mao, Y., Wang, X., and Liu, L. (2022). Enzymological and structural characterization of Arabidopsis thaliana heme oxygenase-1. FEBS Open Bio 12: 1677–1687.

Waters, M.T. and Langdale, J.A. (2009). The making of a chloroplast. EMBO J. 28: 2861–2873.

Xu, F., Jia, M., Li, X., Tang, Y., Jiang, K., Bao, J., and Gu, Y. (2021). Exportin-4 coordinates nuclear shuttling of TOPLESS family transcription corepressors to regulate plant immunity. Plant Cell 33: 697–713.

Yamamoto T., Burke J., Autz G., Jagendorf A.T. (1981). Bound ribosomes of pea chloroplast thylakoid membranes: Location and release in vitro by high salt, puromycin, and RNase. Plant Physiol. 67: 940–949.

Yoo, K.S., Ok, S.H., Jeong, B.C., Jung, K.W., Cui, M.H., Hyoung, S., Lee, M.R., Song, H.K., and Shin, J.S. (2011). Single cystathionine β-synthase domain-containing proteins modulate development by regulating the thioredoxin system in Arabidopsis. Plant Cell 23: 3577–3594.

Zoschke, R. and Bock, R. (2018). Chloroplast translation: Structural and functional organization, operational control, and regulation. Plant Cell 30: 745–770.

Zoschke R., Barkan A. (2015). Genome-wide analysis of thylakoid-bound ribosomes in maize reveals principles of cotranslational targeting to the thylakoid membrane. Proc Natl Acad Sci USA 112: E1678–87.

